## Supplementary figures and images for "MYBL2 regulates ATM to control replication initiation and prevent replication stress in pluripotent stem cells"

### expansion view 1

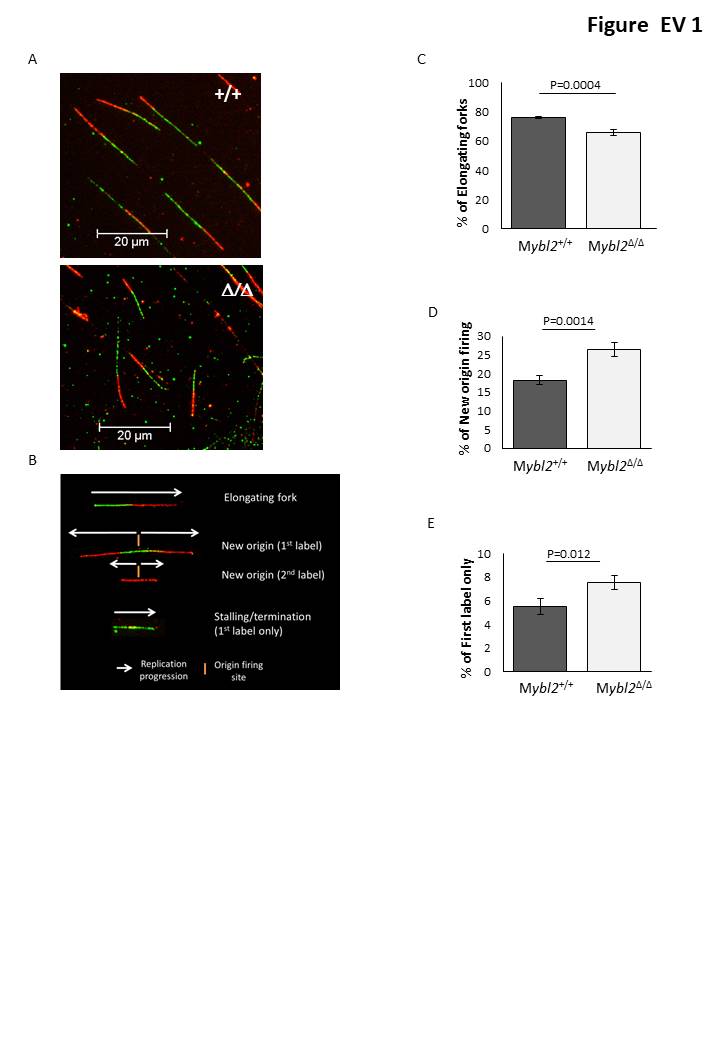

### expansion view 2

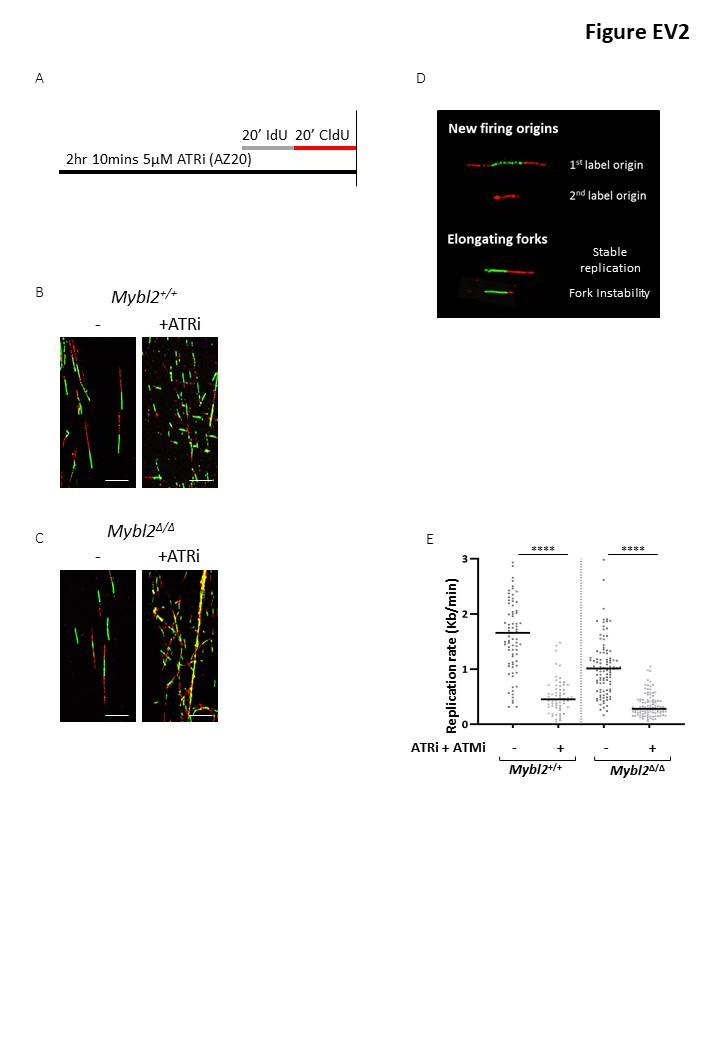

### expansion view 3

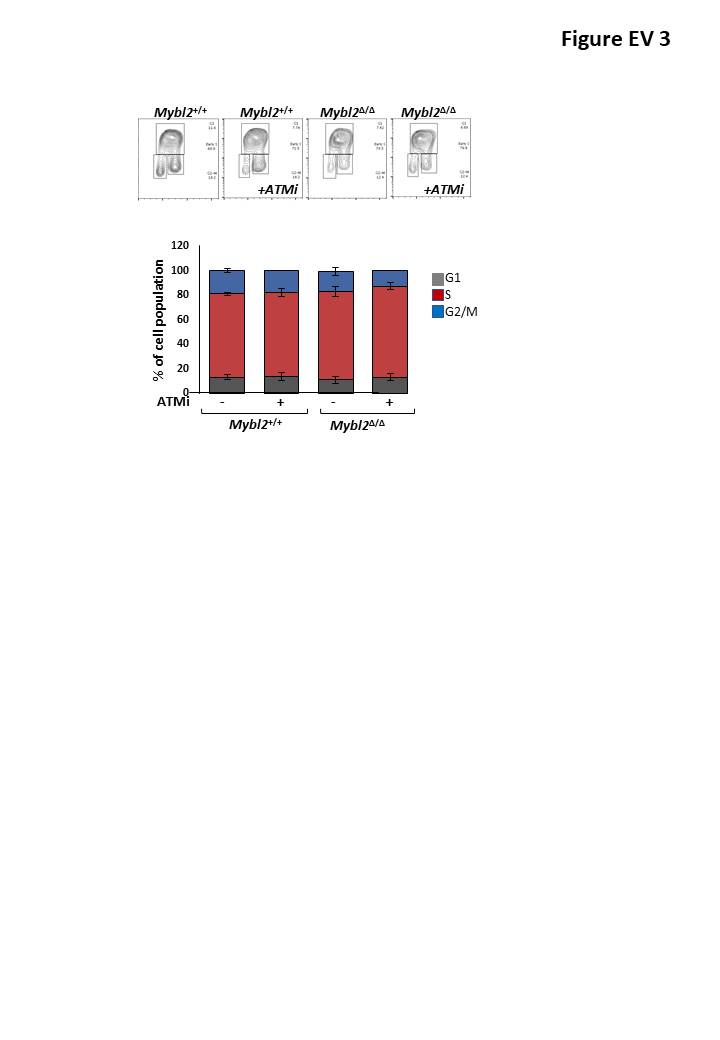

### expansion view 4

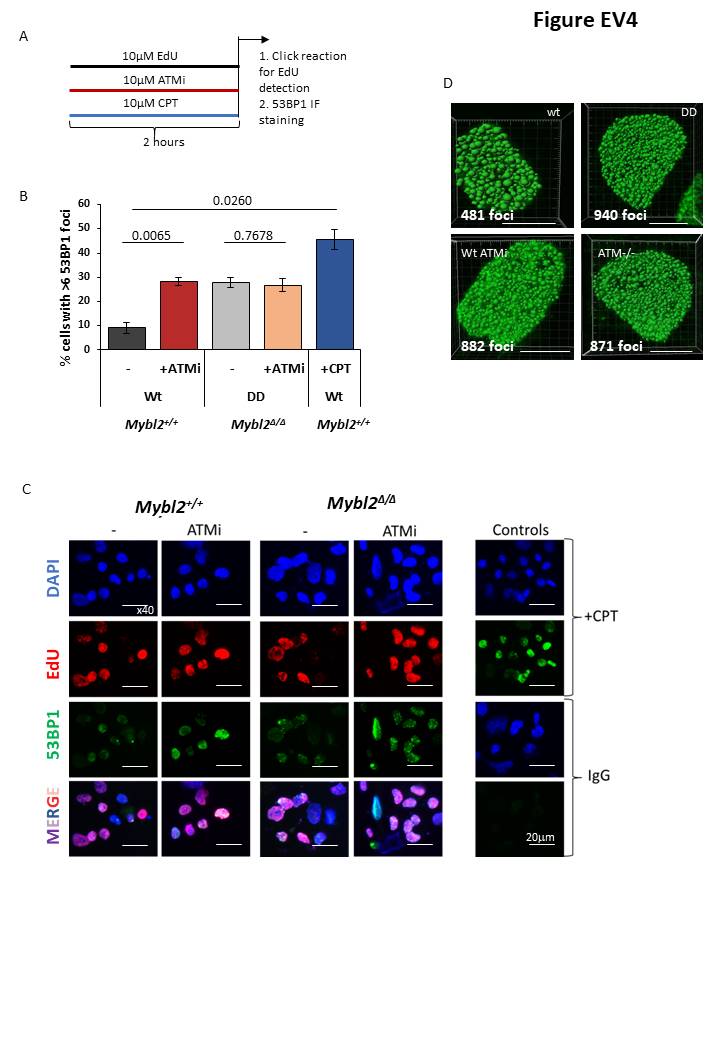

### expansion view 5

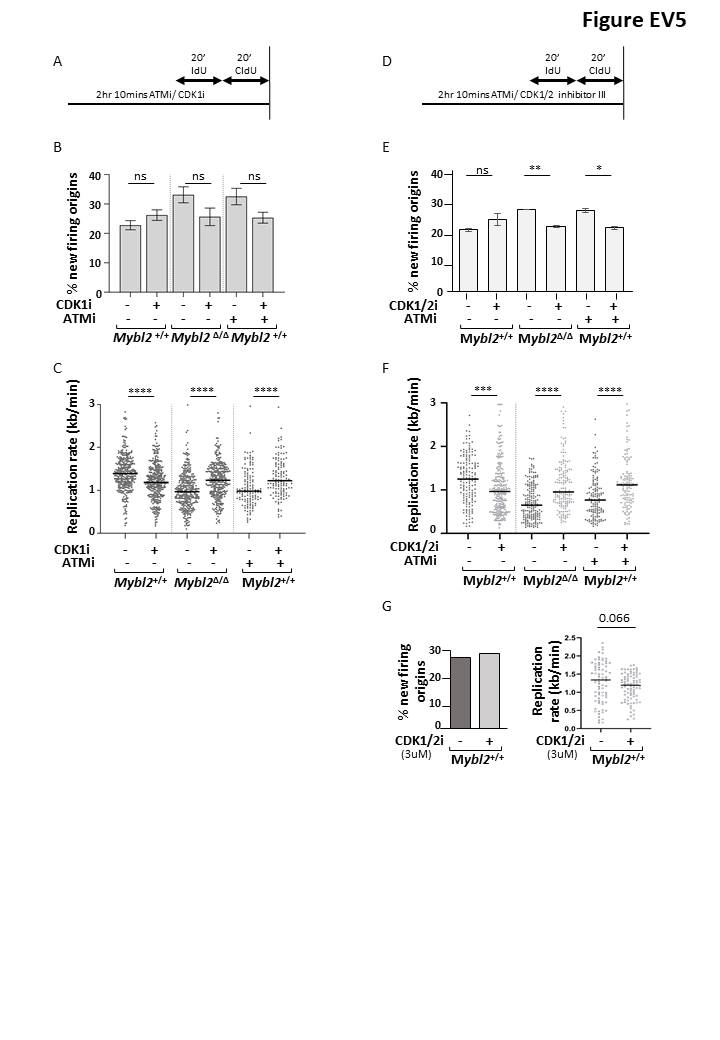
