## Appendix figures for "MYBL2 regulates ATM to control replication initiation and prevent replication stress in pluripotent stem cells"

### Table of context

**Appendix figures**

**Appendix Figure S1. Replicating dependent double strand breaks in *Mybl2*$^{\Delta/\Delta}$ *ESCs*.**

**Appendix Figure S2. Validation of P-CDK1 (Tyr15).**

**Appendix Figure S3. Spontaneous P-CHK1 observed in the untreated *Mybl2*$^{\Delta/\Delta}$ ESCs is ATR-dependent.**

**Appendix Figure S4. ATM regulates replication stress in pluripotent stem cells.**

### Appendix Figure 1

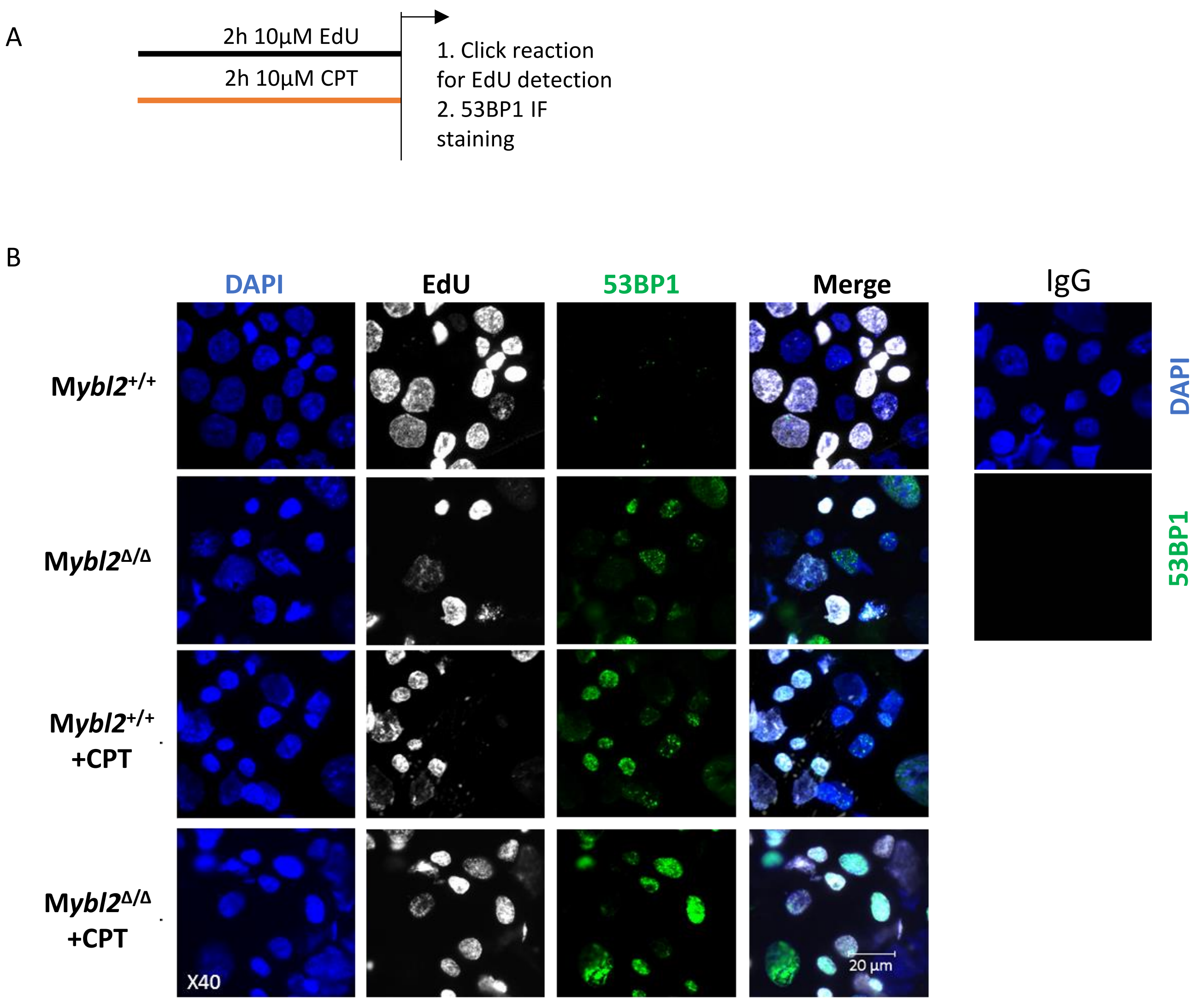


**Appendix Figure S1. Replicating dependent double strand breaks in *$Mybl2^{\Delta/\Delta}$ ESCs*.**
A) Scheme of the protocol. 10µM EdU was incorporated for 2 hours in culture and detected using the 'click' reaction. For a measure of double strand breaks, immunofluorescence staining for 53BP1 was performed. Cells were also treated for 2 hours with 10µM CPT as positive control of damage.
B) Representative images for EdU and 53BP1 immunostaining for *$Mybl2^{+/+}$*, B- *$myb^{\Delta/\Delta}$* ESCs treated or not with CPT. 200 EdU positive were analyzed for 4 experimental repeats.

#### Appendix Figure 2

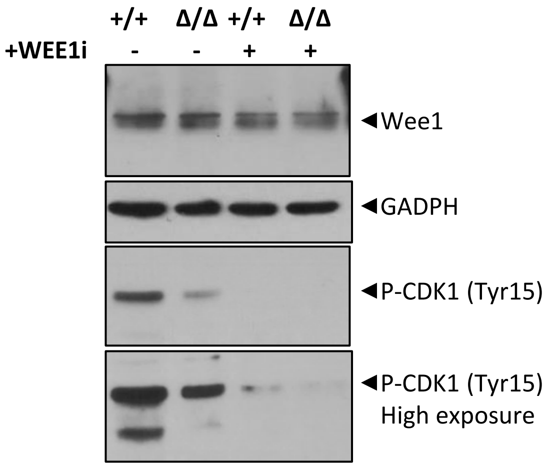


**Appendix Figure S2. Validation of P-CDK1 (Tyr15).**
Wee1, P-CDK1 (Tyr15) and GADPH expression levels of *Mybl2*$^{+/+}$ and *Mybl2*$^{\Delta/\Delta}$ ESCs with or without wee-1 inhibitor treatment analyzed by immunoblotting.

### Appendix Figure S3

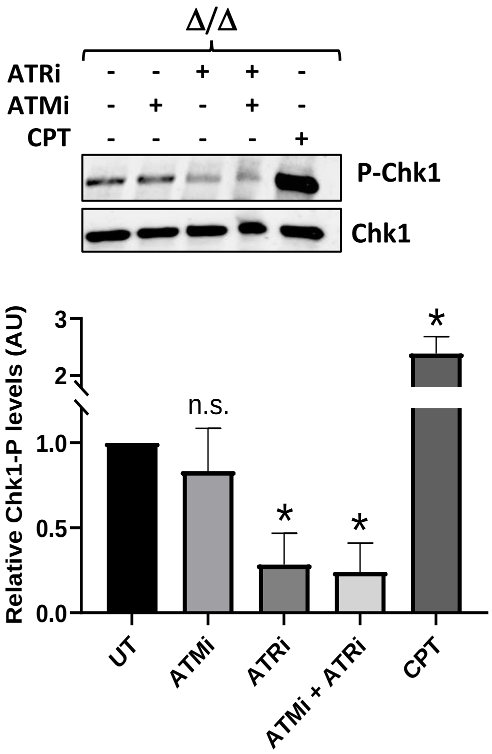


**Appendix Figure S3. Spontaneous P-CHK1 observed in the untreated *Mybl2*$^{\Delta/\Delta}$ ESCs is ATR-dependent.** Immunoblot showing the levels of phosphorylated CHK1 (Ser345) and CHK1 in the *Mybl2*$^{+/+}$ and *Mybl2*$^{\Delta/\Delta}$ ESCs with or without ATM inhibitor (KU66019) 10uM for 2hours; ATR inhibitor (AZ20) 5uM for 2 hours and CPT treatment as positive control (10µM CPT for 2 hours).

### Appendix Figure S4

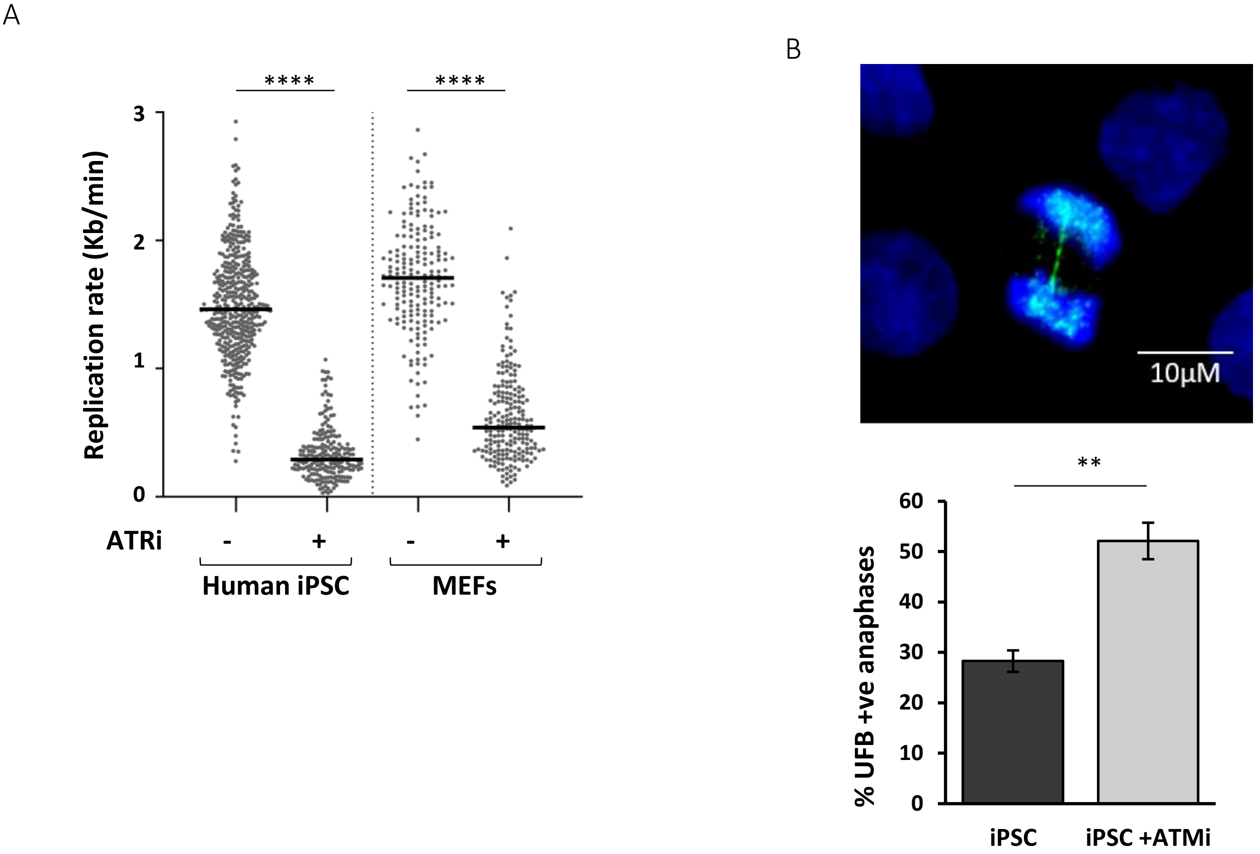


**Appendix Figure S4. ATM regulates replication stress in pluripotent stem cells.**
A) Distribution of replication fork rates for MEFs and human iPSC treated and untreated with ATR inhibitor AZ20. Statistical analysis was performed using the Mann Whitney U test. n=3 experimental replicates. A minimum of 190 replication forks were counted MEFs and a minimum of 220 replication forks for human iPSC.
B) Representative image of UFB in human iPSC treated with ATM inhibitor 10µM KU60019. PITCH protein in Green and DAPI staining in blue.
C) Percentage of anaphases positive for UFB (PITCH positive staining) for human iPSC treated or not with 10µM KU60019. Data represents at least 100 anaphases from each group from 3 experimental repeats.
